# Deep Learning for Non-Invasive Cortical Potential Imaging

**DOI:** 10.1101/2020.06.15.151480

**Authors:** Alexandra Razorenova, Nikolay Yavich, Mikhail Malovichko, Maxim Fedorov, Nikolay Koshev, Dmitry V. Dylov

## Abstract

Electroencephalography (EEG) is a well-established non-invasive technique to measure the brain activity, albeit with a limited spatial resolution. Variations in electric conductivity between different tissues distort the electric fields generated by cortical sources, resulting in smeared potential measurements on the scalp. One needs to solve an ill-posed inverse problem to recover the original neural activity. In this article, we present a generic method of recovering the cortical potentials from the EEG measurement by introducing a new inverse-problem solver based on deep Convolutional Neural Networks (CNN) in paired (U-Net) and unpaired (DualGAN) configurations. The solvers were trained on synthetic EEG-ECoG pairs that were generated using a head conductivity model computed using the Finite Element Method (FEM). These solvers are the first of their kind, that provide robust translation of EEG data to the cortex surface using deep learning. Providing a fast and accurate interpretation of the tracked EEG signal, our approach promises a boost to the spatial resolution of the future EEG devices.

## 1 Introduction

Electroencephalography (EEG) is a common method of non-invasive registration of the brain activity. It has high temporal resolution and has low operational cost. These aspects make the adoption of EEG widespread, including the areas of neurophysiological and cognitive research, non-invasive Brain-Computer Interface (BCI) devices, clinical studies, and diagnostics. However, there are multiple technical limitations to modern EEG technology, with a poor spatial resolution being the major one [20, 28]. The low spatial resolution of EEG is primarily caused by the significant difference of conductivity between the skull and the other tissues. Generally speaking, the conductivity of all tissues composing the head should be taken into account.

In this work, we will consider how to recover the signal from the brain surface given measurements on the scalp, using the most recent arsenal of techniques from the classical and the deep learning disciplines. We proceed with a formal problem statement and with a review of the state-of-the-art.

### 1.1 Statement of the problem

Let the head be represented by the computational domain *Ω* ⊂ ℝ^3^ bounded by a piecewise-smooth boundary *∂Ω*. Within the domain, the electric potential *p*(**x**) satisfies the following expression [6]:

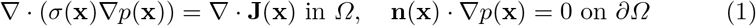

where *σ*(**x**) is the conductivity distribution over the head volume, **J**(**x**) represents the volumetric distribution of a primary current source that produces the signal, and **n**(**x**) is the vector normal to the surface *∂Ω* at the point **x** ∈ *∂Ω*. The conductivity *σ* is assumed to be known. We also introduce the area of measurements on the scalp *Γ* ⊂ *∂Ω*, and the cortex *C* ⊂ *Ω*. Based on equations (1), we can formulate three main mathematical problems (see Fig. 1A for visualizing the concept):

**Fig. 1.**
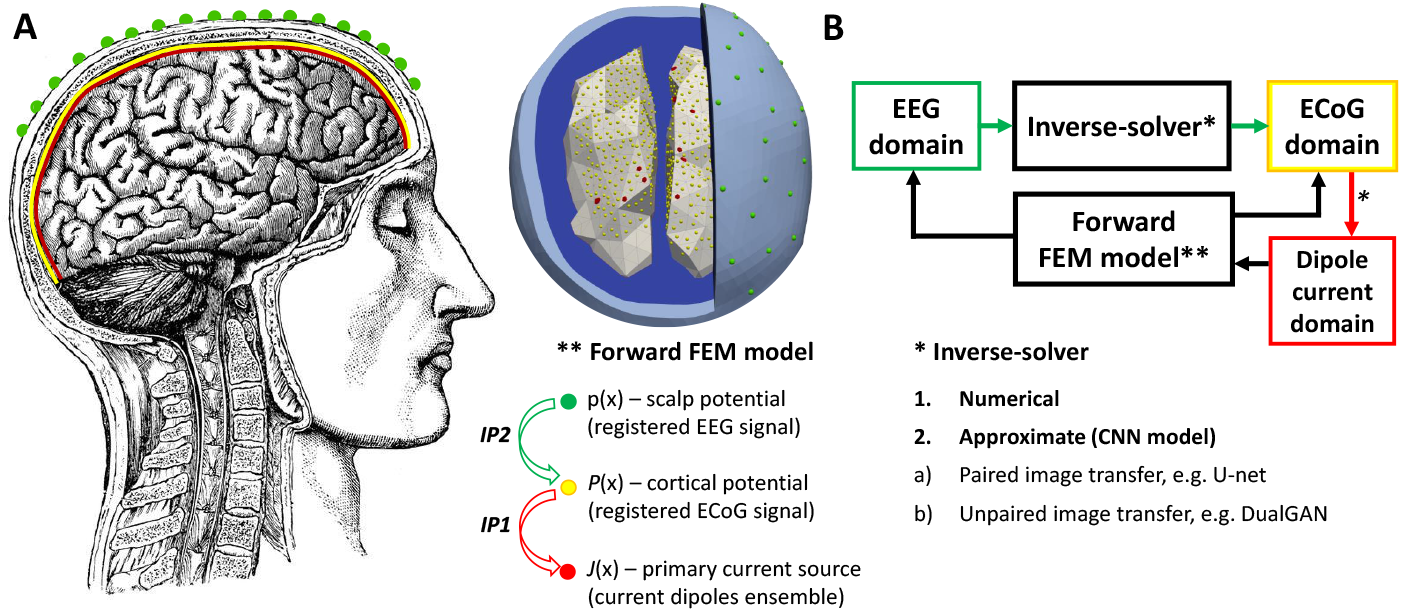
A. Concept of cortical activity registration (left, head image reproduced from [7]) and its physical model implemented in FEM (right). The IP stands for “Inverse problem”. B. Schematic representation of Forward and Inverse problems solving.

**Forward problem (FP):** *compute the scalp voltages p*(**x**), **x** ∈ *∂Ω, given current dipoles or density* **J**(**x**).

**Inverse problem 1 (IP1):** *given known electric voltages p*(**x**), **x** ∈ *Γ on the area of measurements Γ, compute current density* **J**(**x**), **x** ∈ *C on the cortex.*

**Inverse problem 2 (IP2):** *given known electric voltages p*(**x**), **x** ∈ *Γ on the area of measurements Γ, compute the electric voltage p*(**x**), **x** ∈ *C on the cortex.*

### 1.2 State-of-the-art

The majority of algorithms proposed in the EEG community are dedicated to IP1 and formulated in the framework of Tikhonov regularization, including the MNE and LORETA families, etc. (see [12, 24, 23] for a review). The main limitation of these methods is their low spatial resolution due to the head model geometry and the high demand for computational resources (especially, if MRI-based head models are used). There are algorithms for solving IP2 [4, 10, 15] but their spatial resolution is insufficient.

Recent studies propose neural networks as alternative method of the inverse problem solution [14, 29]. Convolutional neural networks (CNN) are capable of providing both an approximation of the physical model and a sufficient regularization, leading to more accurate inverse-problem solution, stable with respect to noisy inputs. U-Net can serve as an efficient integrator of various numerical modelling solutions to improve the accuracy [19]. An Autoencoder-based CNN was also proposed for EEG super-resolution as a non-linear interpolation method [17]. Finally, the most popular trend today is to use the temporal information as an additional constraint and to apply Markov models, or their approximations, in a recurrent network configuration (e.g. LSTM) [18, 13, 5].

In this work, we aspire to solve IP2 using deep CNNs. We present a generic methodology for recovering cortical potentials (referred here as extended ECoG) from the EEG measurement using a new CNN-based inverse-problem solver based on the U-Net or the DualGAN architectures (Fig. 1). Paired examples of EEG-ECoG synthetic data were generated via forward problem solving Eq. (1). Thus, we reformulate the problem as an image-to-image translation task and find a way to reconstruct accurate mapping of the cortical potentials to recover the original brain activity beyond the spatial resolution of EEG.

## 2 Methods

### Head Model and Data Generation

A realistic 3-D head model was used to prepare a synthetic dataset (Fig. 1, A). An anatomical head model was constructed from the sample subject data of the `MNE` package [11]. We extracted triangulated surfaces of the cortex, skull and skin and smoothed them using the `iso2mesh` toolbox [26]. The volume conductor was meshed into 261,565 tetrahedrons with 42,684 vertices. The conductivity of the cortex surface, the skin, and the skull were assumed to be 0.33 S/m, 0.33 S/m, and 0.01 S/m, respectively [9].

Source current dipoles were positioned in the centers of the boundary triangles of the cortex mesh. For simplicity, we considered only the upper part of the cortex in order to provide an 2-D representation of the data, resulting in 400 possible locations of the source current dipoles (the number of active dipoles was, however, restricted for each simulation to *n* dipoles, as described in section 3). Cortical potential sensors were located in the same manner, leading to 400 measurement probes of the cortex potential (see yellow dots in Fig. 1A, left). Scalp potential sensors (i.e., channels of an EEG device) were uniformly located on the outer scalp surface. We tested different number of EEG channels: 128, 64, or 32.

Our simulation run went as follows. First, a random initialization of *n* current dipoles with the current values in the range 0.1-0.9 *μ*A [22] was done. Then, we carried out a calculation of EEG and ECoG data as a numerical solution to the FP (1), discretized with the Finite Element Method (FEM) [2]. The resulting system was solved with preconditioned conjugate gradient method [8]. Thus, EEG and the ECoG pairs were modelled. We converted them into 256 × 256 float32 images to provide precise topographic representation of the measured activity. Min-max contrast normalization was applied to the image intensities.

Thus, each computed output is a pair of the top-view image of the scalp potential distribution and the top view image of the cortical potential distribution.

### Evaluation details

Using the head model, a dataset of synthetic EEG-ECoG pairs was generated. It was split into 5000 train, 400 validation, and 600 test pairs. Both the data generation and the training of the neural network were performed on the [Anonimized] supercomputer using Tesla V100-SXM2 GPUs [1].

To train the topographic translation from EEG to ECoG domain, we did minimal modifications to U-Net [27] (see Fig. 2) to adapt it to the paired image-to-image translation task. The U-Net network was optimized to minimize the Binary Cross Entropy (BCE) loss function between the recovered ECoG image and the corresponding ground truth (GT) ECoG image. The training was performed with the ADAM optimizer, sigmoid activation function, the batch size of 4, and the learning rate of 10^-4^.

**Fig. 2.**
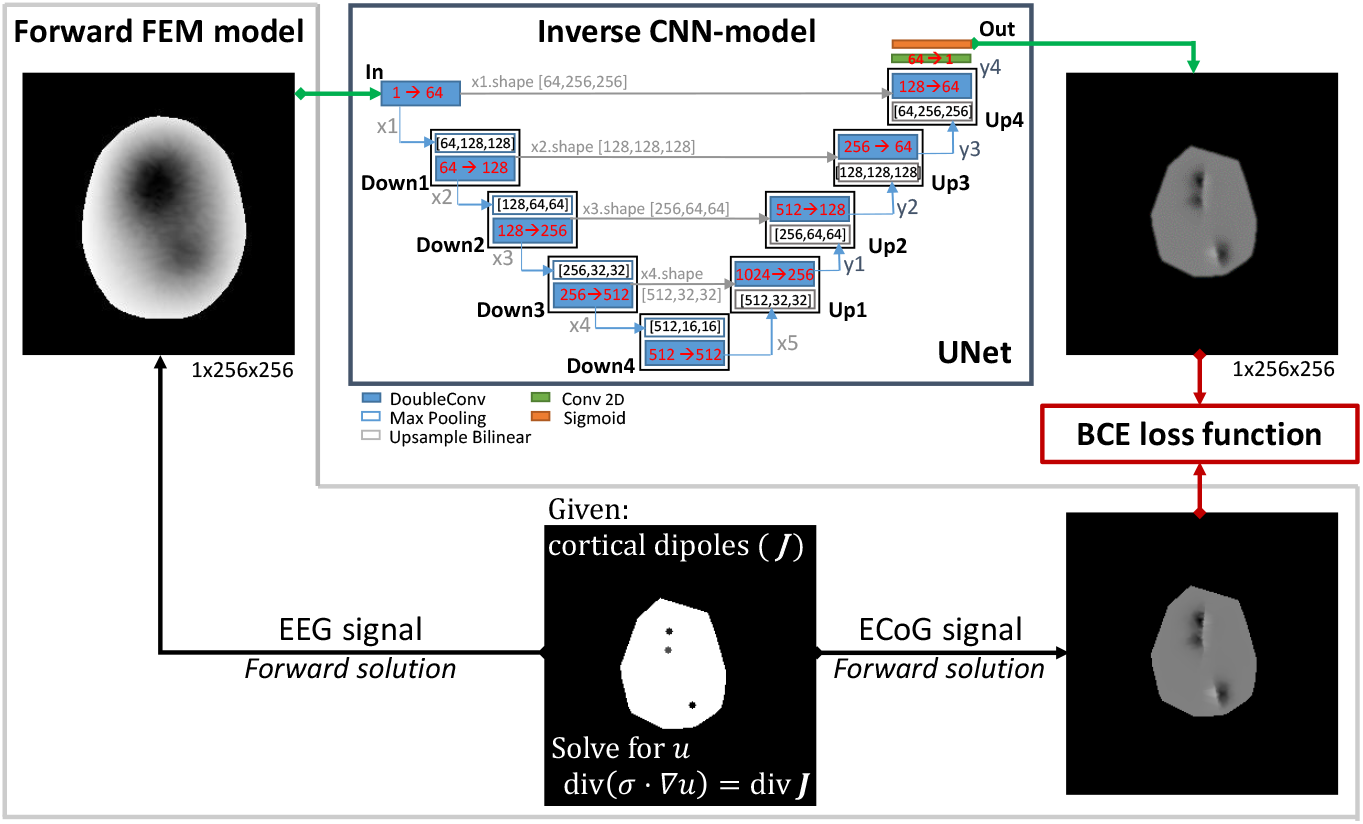
U-Net-solver for imaging cortical potential from the EEG data. The EEG and the ECoG images are modelled via solution to the FP, defined in text.

Additionally, we tested DualGAN architecture [30, 25] as an inverse solver, aiming at creating an architecture to handle realistic unpaired EEG-ECoG data samples available in clinical practice, e.g. [3]. Its first generator was trained to translate ECoG images to EEG; and the second generator translated the images from EEG to ECoG domain.

## 3 Experiments

We tested the effectiveness and robustness of the proposed U-Net-solver by varying the complexity of the pattern of cortical activity. For this purpose, the proposed model was trained on four datasets with different number of active dipoles.

We started from a subset where the corresponding ECoG and the EEG image representations were generated from a superposition of one, two, or three source current dipoles. Then, we gradually increased the model’s complexity to 5-10, 10-25, and 25-45 active dipoles, respectively (see Fig. 3, top row).

**Fig. 3.**
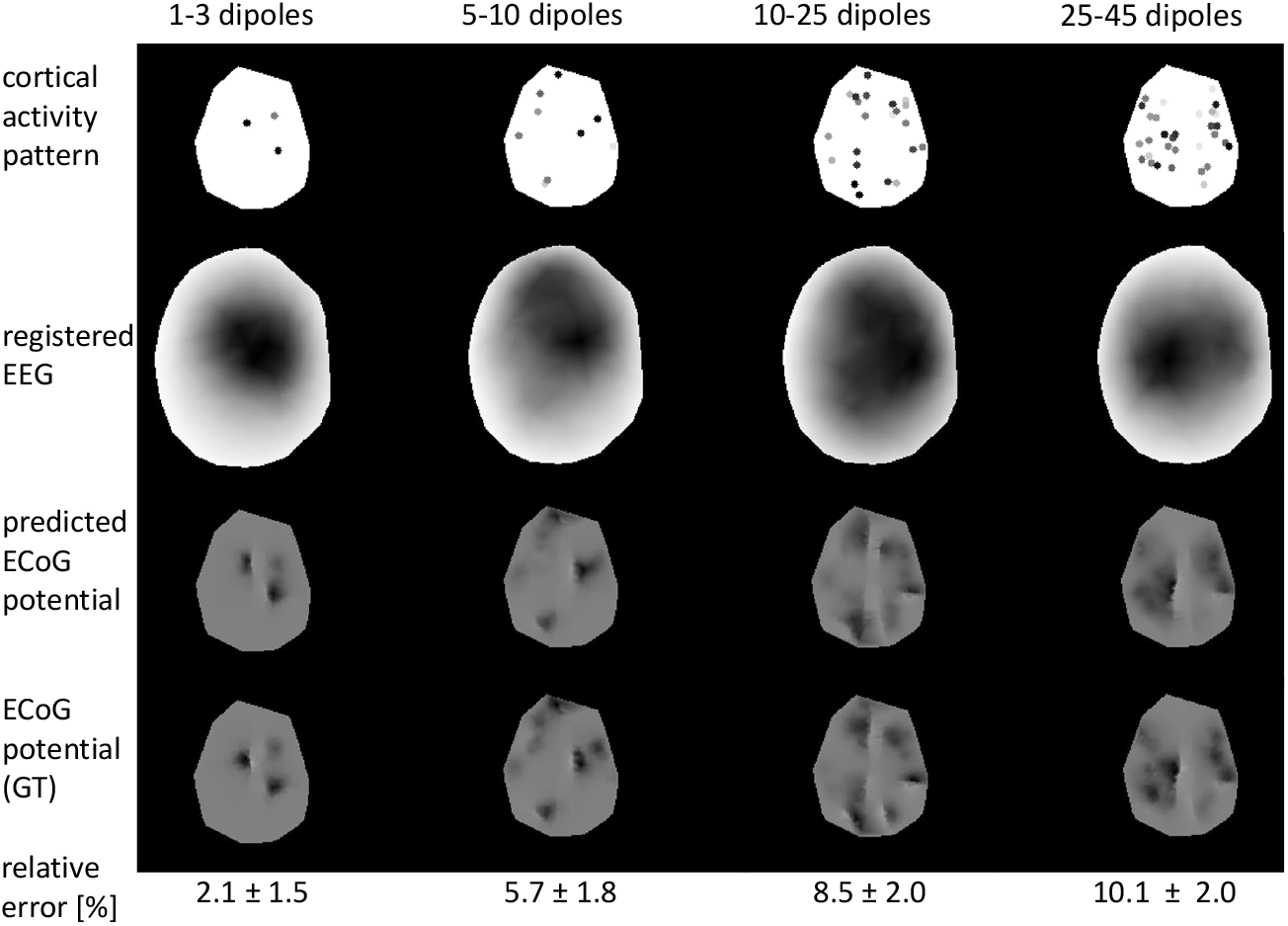
The U-Net-solver results for cortical potential imaging as a function of a random number of randomly located source dipoles (columns from left to right correspond to the increased number of sources).

As can be seen in the Fig. 3, the U-Net-solver is capable of recovering the original cortical potential distribution pattern given the indiscernible EEG data input. The relative error between the ground truth cortical potential distribution and the one obtained via the U-Net-solver is lower than 10%. The error does not increased dramatically as the complexity of the original cortical activity pattern is varied.

We noted that the relative error dynamics can be explained by the number of dipoles and by the variability of this parameter within the training set. In other words, when 1 to 3 dipoles are active simultaneously, the majority of the training examples is separable and the reconstruction of the sources lacks any ambiguity, but when the number of active dipoles rises to 10 to 25 dipoles, the separation of the source dipoles on the ground truth images is less pronounced. This additional ambiguity limits the overfitting of the solver.

To study the generalization performance of the U-Net-solver all trained models were reciprocally tested on all test sets. The experiments were done for the 128-channel EEG. We also tested the ability of the solver to process the EEG data, obtained from more commonly used EEG configurations: 64 and 32 channels. For this experiment, pattern of only 5-10 dipoles were considered, and the result is shown in Fig. 4 (inset table).

**Fig. 4.**
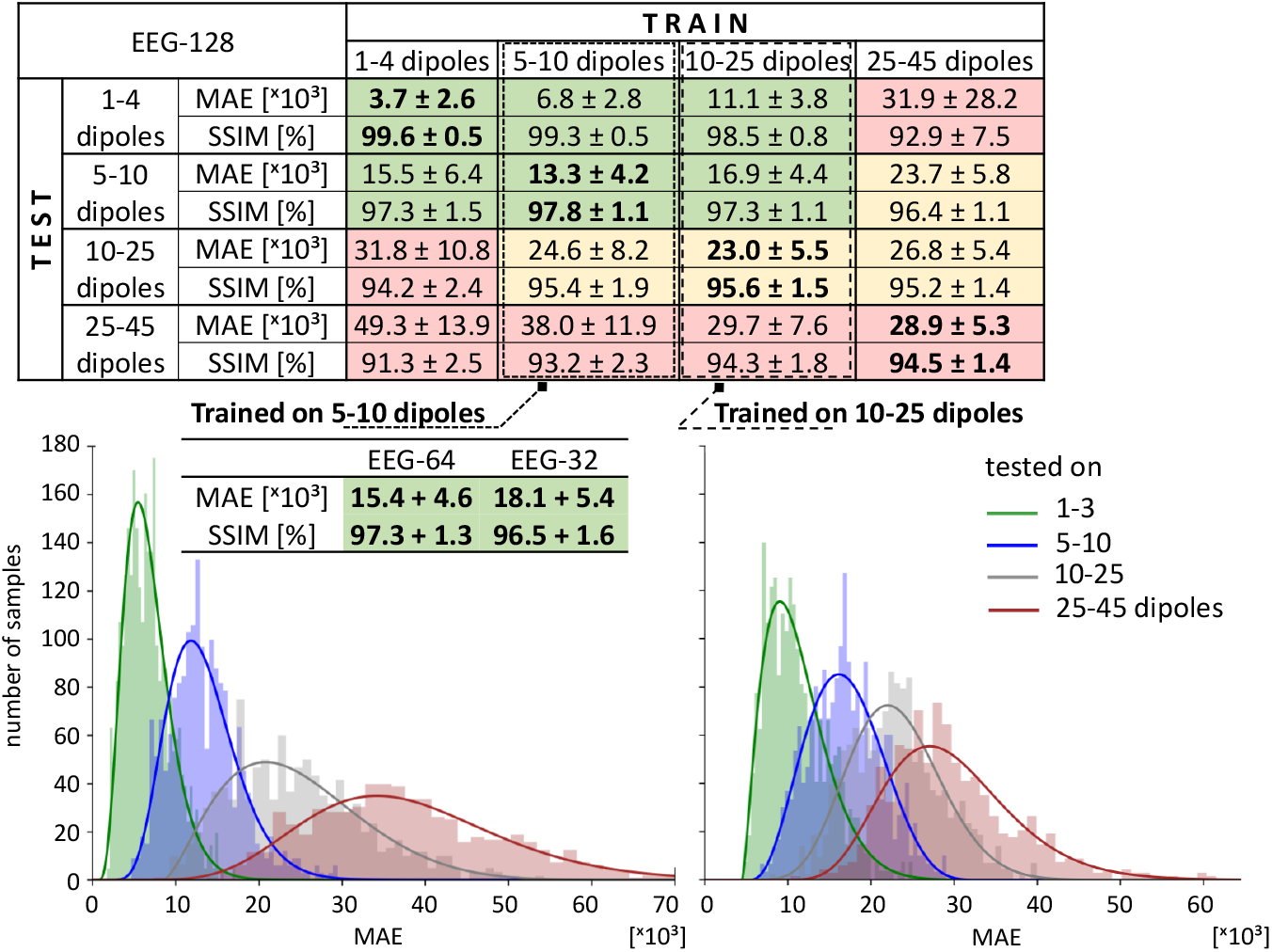
Performance of the U-Net-solver as a function of a number of active dipoles and a number of EEG channels. Cells, where both the train and the test data were generated with the same number of dipoles and the same number of channels, are marked in bold. Histograms on the bottom show MAE distribution for the data used as the test sample; the models were trained with 5-10 (left) and 10-25 (right) dipoles.

Conventional metrics, such as MAE and SSIM, were calculated for ECoG topomaps. Background pixels were excluded. All cells in the tables of Fig. 4 are color-coded using empirical threshold: cells with relatively high performance (MAE*<*20 c.u. and SSIM*>*95%) are highlighted in green; cells with relatively moderate performance (MAE*<*30 c.u. or SSIM*>*95%) are highlighted in yellow; the other cells are highlighted in red.

We observe that the U-Net-solver trained on 10-25 dipoles yields similar errors for pairs containing 1-3 or 5-10 dipoles and thus is capable of generalizing the input data. Therefore it effectively resolves the EEG signal even if the configuration differs from the source pattern which the solver was trained on. The trade-off between the solver’s accuracy and its ability to generalize is demonstrated in the histograms (Fig. 4, bottom panel).

The solver has a better ability to process unseen data when it was trained on a dataset with more variability. However, it does not hold for the model trained on more complex data examples: e.g., ECoG patterns from a superposition of 25-45 dipoles are reconstructed more coarsely and the pre-trained solver fails to resolve the simpler data examples. This is possibly caused by the loss of information incurred by inaccurately representing the complex cortical patterns as 2D images instead of 3D volumes. In contrast, the U-Net-solver effectively deals with EEG data in any available data configuration. The reduction of information in the input data recorded with 32 or 64 EEG channels (instead of 128 full channel set) does not significantly alter the efficiency of the U-Net-solver, as seen in the extended column 2 of the table in Fig. 4.

## 4 Discussion

### Baseline comparison

Since to our best knowledge ours is the first DL-based approach, we compared our U-Net-solver to the numerical solution of the Cauchy problem by the method of quasi-reversibility [15], using identical inputs for both solvers (see Fig. 5A).

**Fig. 5.**
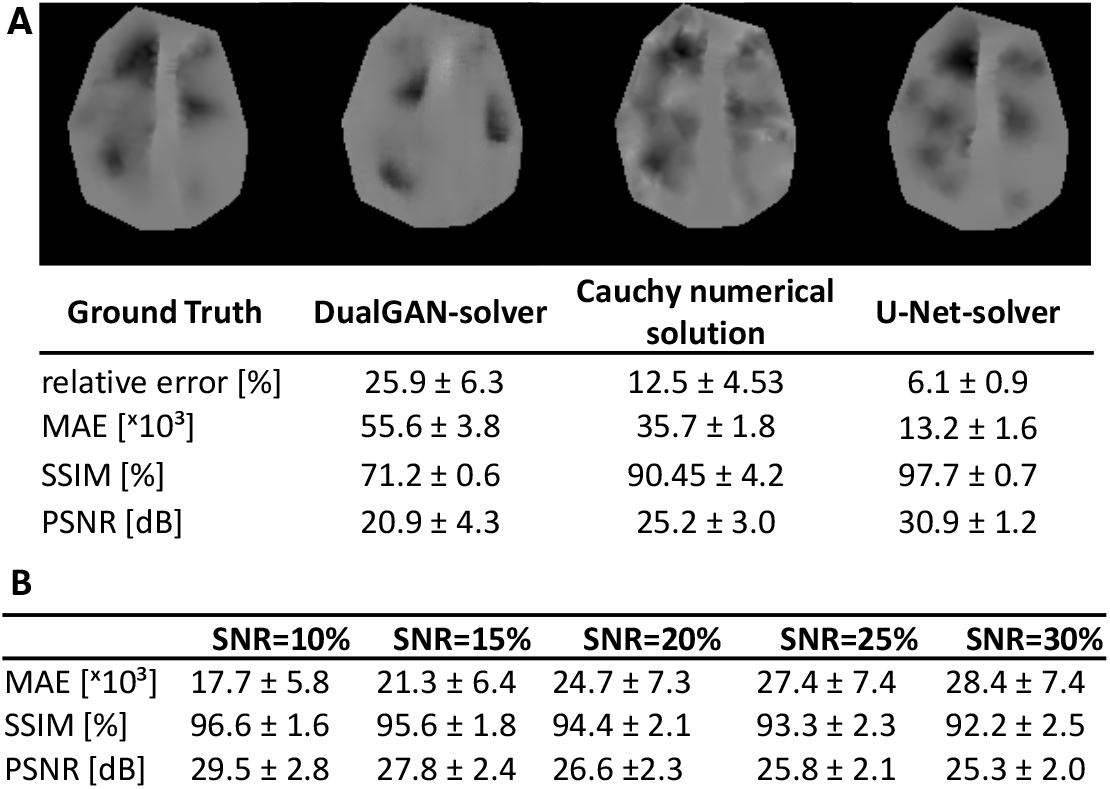
**A.** Comparison of our U-Net-solver to the SOTA method (numerical solution of the Cauchy problem by quasireversibility [15]) and to the DualGAN-solver. **B.** To estimate the stability of U-Net-solver we gradually increased the noise and estimated the error and the image quality scores.

Specifically, the same set of 128 EEG-channel data with 5% Gaussian noise was taken as input images for the U-Net-solver, the DualGAN-solver, and the numerical solution to the Cauchy problem. The statistics calculated on 25 test samples (5-10 dipoles superposition) is shown in the table insets of Fig. 5A.

Once fully trained, our deep learning models exhibit better suitability for the future cortical potential imagers than the numerical method [15]. Their inherent numerical stability shows better results quantitatively (see the scores in Fig. 5) and qualitatively (notice blur patterns in the images). Unlike direct numerical solutions to the ill-posed problem, our models require no explicit regularization to stabilize the approximating functions, avoiding the buildup of error, that is frequent to a particular numerical method. The choice of regularization coefficients constitutes a separate engineering problem that we eliminate herein. Our solvers rely on neural networks that provide implicit regularization of the solution and, ultimately, yield superior accuracy with reduced regularization noise. We also observe that the quality of the solution decreases linearly as the data noise is increased (Fig. 5B), suggesting that our solution is stable with respect to SNR.

Although the U-Net-solver is found to be a better model for reconstructing unseen ECoG patterns, it requires *paired* EEG-ECoG datasets. DualGAN model, to the contrary, could be trained in an *unpaired* manner (Fig. 5A), which increases the amount of readily available data. Preliminary experiments have shown promising results as to the appearence of ECoG topomaps obtained with DualGAN. However, quantitative evaluation demonstrated that the DualGAN architecture cannot be used “as is”. The modification of DualGAN for the optimization of MAE, SSIM and PSNR will be the subject of future research. Another area of further research is the extension of our approach to the translation between corresponding 3-D volumes [21], thus avoiding the loss of information caused by the in-plane projections of the measurements. Such synthetic models can help to pre-train actual clinical translation models embedded into imaging systems, similarly to modern virtual brain models (see, e.g., [16] and references therein).

## 5 Conclusions

We proposed a new approach to brain activity reconstruction by reformulating it into an image-to-image translation framework. We demonstrated successful 2-D topographical mapping of the EEG data to the cortical potentials. CNNs fine-tuned to directly approximate the inverse-operator, achieved a remarkably low error of reconstruction and high image quality scores, effectively promising a boost to the spatial resolution of the EEG signal. The CNN-based solver demonstrated high stability with respect to noise and to the number of measurements (EEG channels). Due to the flexibility of the solver, it can provide approximations to the solution regardless of the geometries of the scalp and the cortex, mitigating possible anatomical differences in patients. Furthermore, the U-Net-solver approach provides accurate primary current dipole localization either with discrete or with distributed source current activity (see Supplementary materials). Our framework can be put at use in areas where high-resolution and high-speed EEG data interpretation is sought after, e.g., in the non-invasive brain-computer interface devices or in localization of epileptogenic foci.

## Notes

### Competing Interest Statement

The authors have declared no competing interest.

## References

1. Anonymized

2. Anderson, R., Andrej, J., Barker, A., Bramwell, J., Camier, J.S., Cerveny, J., Dobrev, V., Dudouit, Y., Fisher, A., Kolev, T., et al.: Mfem: a modular finite element methods library. arXiv preprint 1911.09220 (2019)

3. Boran, E., Fedele, T., Steiner, A., Hilfiker, P., Stieglitz, L., Grunwald, T., Sarnthein, J.: Dataset of human medial temporal lobe neurons, scalp and intracranial eeg during a verbal working memory task. Scientific Data 7(1), 1–7 (2020)

4. Bourgeois, L.: Convergence rates for the quasi-reversibility method to solve the Cauchy problem for Laplace’s equation. Inverse problems 22(2), 413 (2006)

5. Cui, S., Duan, L., Gong, B., Qiao, Y., Xu, F., Chen, J., Wang, C.: EEG source localization using spatio-temporal neural network. China Communications 16(7), 131–143 (2019)

6. Da Silva, F.L., Hansen, P., Kringelbach, M., Salmelin, R., et al.: Electrophysiological basis of meg signals. In: MEG: an introduction to methods, pp. 1–2. Oxford Univ. Press (2010)

7. Fischer, A., Kaplan, M., Azéma, L.: La femme, Médecin du Foyer: Ouvrage d’Hygiène et de Médecine familiale, concernant particulièrement les Maladies des Femmes et des Enfants, les Accouchements et les Soins a’ donner aux Enfants. E. Posselt & Cie, Éditeurs, Bibliothèque nationale de France (1905), http://catalogue.bnf.fr/ark:/12148/cb30437824k

8. Fletcher, R.: Conjugate gradient methods for indefinite systems. In: Numerical analysis, pp. 73–89. Springer (1976)

9. Fuchs, M., Wagner, M., Kastner, J.: Development of volume conductor and source models to localize epileptic foci. Journal of Clinical Neurophysiology 24(2), 101–119 (2007)

10. Gevins, A., Le, J., Brickett, P., Reutter, B., Desmond, J.: Seeing through the skull: advanced EEGs use MRIs to accurately measure cortical activity from the scalp. Brain Topography 4(2), 125–131 (1991)

11. Gramfort, A., Luessi, M., Larson, E., Engemann, D., Strohmeier, D., Brodbeck, C., Goj, R., Jas, M., Brooks, T., Parkkonen, L., Hämäläinen, M.: MEG and EEG data analysis with MNE-Python. Frontiers in Neuroscience 7, 267 (2013). https://doi.org/10.3389/fnins.2013.00267, https://www.frontiersin.org/article/10.3389/fnins.2013.00267

12. Hamalainen, M.S., Ilmoniemi, R.J.: Interpreting magnetic fields of the brain: minimum norm estimates. Medical & biological engineering & computing 32(1), 35–42 (1994)

13. Hansen, S.T., Hansen, L.K.: Spatio-temporal reconstruction of brain dynamics from EEG with a Markov prior. NeuroImage 148, 274–283 (2017)

14. Jin, K.H., McCann, M.T., Froustey, E., Unser, M.: Deep convolutional neural network for inverse problems in imaging. IEEE Transactions on Image Processing 26(9), 4509–4522 (2017)

15. Koshev, N., Yavich, N., Malovichko, M., Skidchenko, E., Fedorov, M.: Fem-based scalp-to-cortex eeg data mapping via the solution of the cauchy problem. Journal of Inverse and Ill-posed Problems (0) (2020)

16. Krylov, D., Dylov, D.V., Rosenblum, M.: Reinforcement learning for suppression of collective activity in oscillatory ensembles. Chaos: An Interdisciplinary Journal of Nonlinear Science 30(3), 033126 (2020). https://doi.org/10.1063/1.5128909

17. Kwon, M., Han, S., Kim, K., Jun, S.C.: Super-resolution for improving EEG spatial resolution using deep convolutional neural network—feasibility study. Sensors 19(23), 5317 (2019)

18. Lamus, C., Hämalainen, M.S., Temereanca, S., Brown, E.N., Purdon, P.L.: A spatiotemporal dynamic distributed solution to the MEG inverse problem. NeuroImage 63(2), 894–909 (2012)

19. Latvala, J.: Applying neural networks for improving the MEG inverse solution. G2 pro gradu, diplomityö (2017-12-11), http://urn.fi/URN:NBN:fi:aalto-201712188173

20. Lopes da Silva, F.: EEG and MEG: Relevance to neuroscience. Neuron 80(5), 1112–1128 (2013). https://doi.org/https://doi.org/10.1016/j.neuron.2013.10.017,http://www.sciencedirect.com/science/article/pii/S0896627313009203

21. Milletari, F., Navab, N., Ahmadi, S.A.: V-net: Fully convolutional neural networks for volumetric medical image segmentation. In: 2016 Fourth International Conference on 3D Vision (3DV). pp. 565–571. IEEE (2016)

22. Murakami, S., Okada, Y.: Contributions of principal neocortical neurons to magnetoencephalography and electroencephalography signals. The Journal of physiology 575(3), 925–936 (2006)

23. Pascual-Marqui, R., Sekihara, K., Brandeis, D., Michel, C., Koenig, T., Gianotti, L., Wackermann, J.: Imaging the electric neuronal generators of EEG/MEG. Electrical Neuroimaging p. 49–78 (2009)

24. Pascual-Marqui, R.D., et al.: Standardized low-resolution brain electromagnetic tomography (sLORETA): technical details. Methods Find Exp Clin Pharmacol 24(Suppl D), 5–12 (2002)

25. Prokopenko, D., Stadelmann, J.V., Schulz, H., Renisch, S., Dylov, D.V.: Unpaired synthetic image generation in radiology using gans. In: Nguyen, D., Xing, L., Jiang, S. (eds.) Artificial Intelligence in Radiation Therapy. pp. 94–101. Springer International Publishing, Cham (2019)

26. Qianqian Fang, Boas, D.A.: Tetrahedral mesh generation from volumetric binary and grayscale images. In: 2009 IEEE International Symposium on Biomedical Imaging: From Nano to Macro. pp. 1142–1145 (June 2009). https://doi.org/10.1109/ISBI.2009.5193259

27. Ronneberger, O., Fischer, P., Brox, T.: U-net: Convolutional networks for biomedical image segmentation. In: International Conference on Medical image computing and computer-assisted intervention. pp. 234–241. Springer (2015)

28. Ryynanen, O.R.M., Hyttinen, J.A.K., Malmivuo, J.A.: Effect of measurement noise and electrode density on the spatial resolution of cortical potential distribution with different resistivity values for the skull. IEEE Transactions on Biomedical Engineering 53(9), 1851–1858 (Sep 2006). https://doi.org/10.1109/TBME.2006.873744

29. Stadelmann, J., Schulz, H., van der Heide, U., Renisch, S.: Pseudo-CT image generation from mDixon MRI images using fully convolutional neural networks. In: Medical Imaging 2019: Biomedical Applications in Molecular, Structural, and Functional Imaging. vol. 10953, p. 109530Z. International Society for Optics and Photonics (2019)

30. Yi, Z., Zhang, H., Tan, P., Gong, M.: DualGAN: Unsupervised dual learning for image-to-image translation. In: Proceedings of the IEEE international conference on computer vision. pp. 2849–2857 (2017)

