## Supplementary materials for "Deep Learning for Non-Invasive Cortical Potential Imaging" (from another version of this preprint: version 1, 2020-06-15)

### Deep Learning for Cortical Potential Imaging – Supplementary material

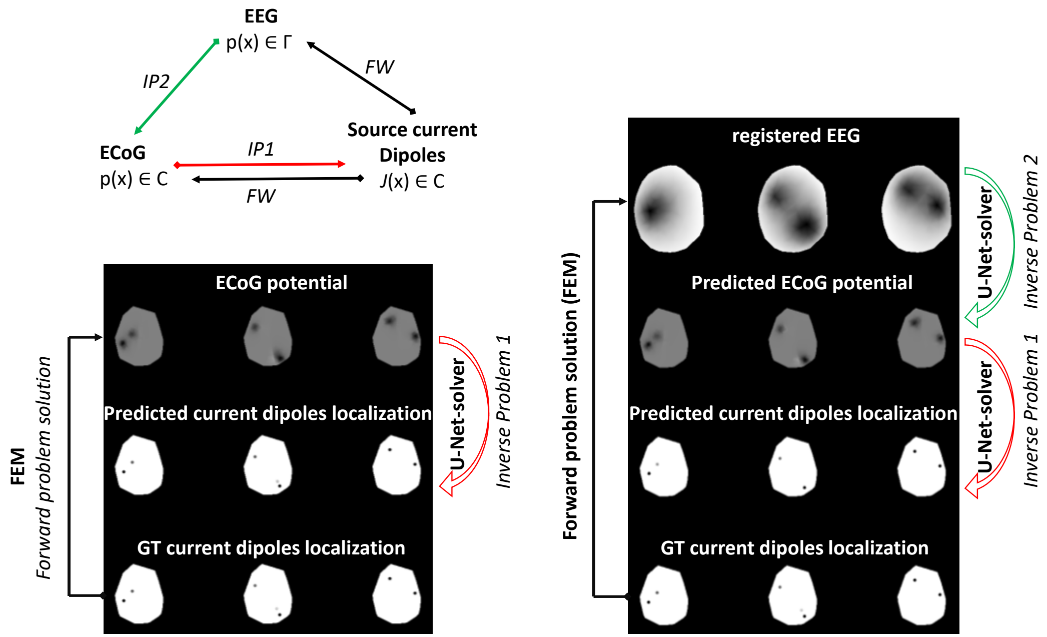


**Fig. 1.** In order to solve IP1, we generated both EEG and ECoG potential from a given dipole localization and solved the forward problems via FEM (top left). We trained U-Net-solver to predict the dipole localization from the ECoG potentials (red arrow). Notice the small difference between the reference ECoG potentials modelled by FEM and those predicted by the U-Net-solver (green arrow). These differences can be explained by distortions from the scalp, skull and meninges represented in the FEM-model.

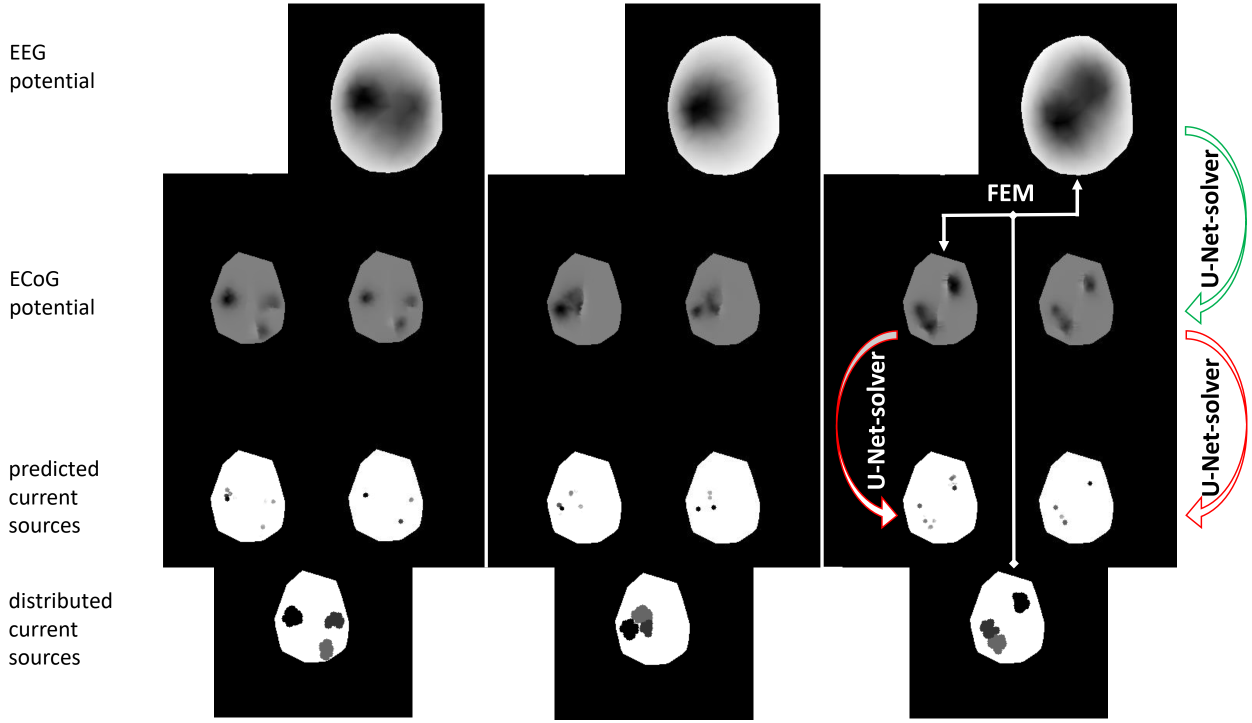


**Fig. 2.** We used FEM to solve the forward problem and generate the EEG and ECoG potentials corresponding to more natural patterns of cortical activity (distributed current sources). The ECoG-to-Source U-Net solver (red arrow) was trained on punctual sources and therefore is not predicting distributed current dipoles. However, the localization of the current sources is accurate: the predicted sources coincide spatially with the reference source pattern. Likewise, the ECoG potentials predicted by the U-Net-solver (green arrow) are similar to their FW-modelled references originated from distributed sources (white arrow) that confirms the generalization capabilities of our EEG-to-ECoG U-Net solver.
